# Measuring plant diversity in a two-stage sampling design by Bayesian updated relative abundances

**DOI:** 10.1101/2022.03.23.485475

**Authors:** Christian Damgaard, Malthe Holst Pedersen, Nikolaj Bjerg Bendsen, Ditte Lønborg Mikkelsen, Bodil K. Ehlers, Thomas Bataillon

## Abstract

The two-stage sampling design provides good local estimates of both the number of plant species and the relative abundances. However, it is a problem to calculate Hill diversity indices at the local scale, because some of the species found in the large plot are not present in the small plot and such species should then incorrectly be weighted with zero relative abundance. A new method for calculating local Hill diversity indices from species richness and relative abundances data is therefore needed. We suggest to replace the local relative abundances with Bayesian updated relative abundance estimates, where the prior probability distribution of the relative abundances are empirically estimated from all plots of the same habitat types. The method is applied on Danish Nardus grasslands.

## Introduction

During the Anthropocene there has been an overall decrease in biodiversity (IPBES 2019). This trend is of general concern, and in order to understand the underlying causes and possibly to reverse this trend, it is important that we have access to unbiased and credible measurements of species diversity.

The most intuitive and simple measure of species diversity in a community is species richness, i.e. the number of species found at a location. However, estimates of species richness are strongly influenced by the presence of rare species that are hard to detect and thus highly sensitive to both sampling effort and relative abundance. Estimates of species richness are highly uncertain and it is often not possible to compare locations because species richness are often estimated using different sampling efforts (Haegeman et al. 2013; Roswell et al. 2021).

Instead of measuring diversity by species richness, it is preferable to use species diversity indices, such as Shannon or Simpson indices, where species occurrence is weighted with its relative abundance in the local community. Moreover, it has been recommended to use the Hill diversity transformation of diversity indices, since they are on the same scale as species richness (Hill 1973; Jost 2006; Roswell et al. 2021).

Hill diversity is generally defined for different weighting functions of the relative abundances, but here we will mainly focus on the Hill-Shannon diversity index, which often is the recommended diversity index (e.g. Roswell et al. 2021), and is defined as,

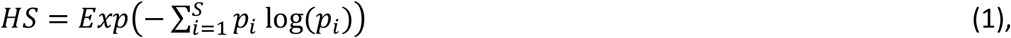

where *S* is the number of species, and *p_i_* is the relative abundance of species *i*.

For calculating Hill diversity indices we need estimates of both the number of species and their relative abundances. Plant abundance is typically estimated non-destructively by measuring plant cover, which is the relative area of a plant species when it is projected onto the soil surface (Damgaard and Irvine 2019). However, it also possible to measure the abundance destructively by harvesting the aboveground biomass and weighing the plant material after it has been sorted into different species. Theoretically, the estimation accuracy and precision of both species richness and relative abundances increases with plot size, and especially so in spatially aggregated plant communities (Kenkel and Podani 1991). However, whereas it is often feasible to get an unbiased estimate of the number of plant species in a relatively large plot, it is often not possible to correctly estimate plant abundance in large plots. Visual estimation of cover in relatively large plots has been shown to be biased and subjective and it only feasible to use the more accurate pin-point method in relatively small plots (see references in Damgaard and Irvine 2019).

Consequently, when measuring Hill diversity indices in a plant community, it will be an advantage to use a two-stage sampling design, which consists of a relatively large plot where the number of species is estimated and a subsample of a relatively small plot within the larger plot, where abundances are estimated. For example, a two-stage sampling design is used in the Danish monitoring program NOVANA, where more than 100.000 plots have been sampled from different habitat types in the period from 2004. In this design (Fig. 1), species richness is estimated in circles with 5 meter radius, and at the center of this circle plant cover data was estimated in 0.5m x 0.5m quadrates using the pin-point method (Nielsen et al. 2012).

**Fig. 1.**
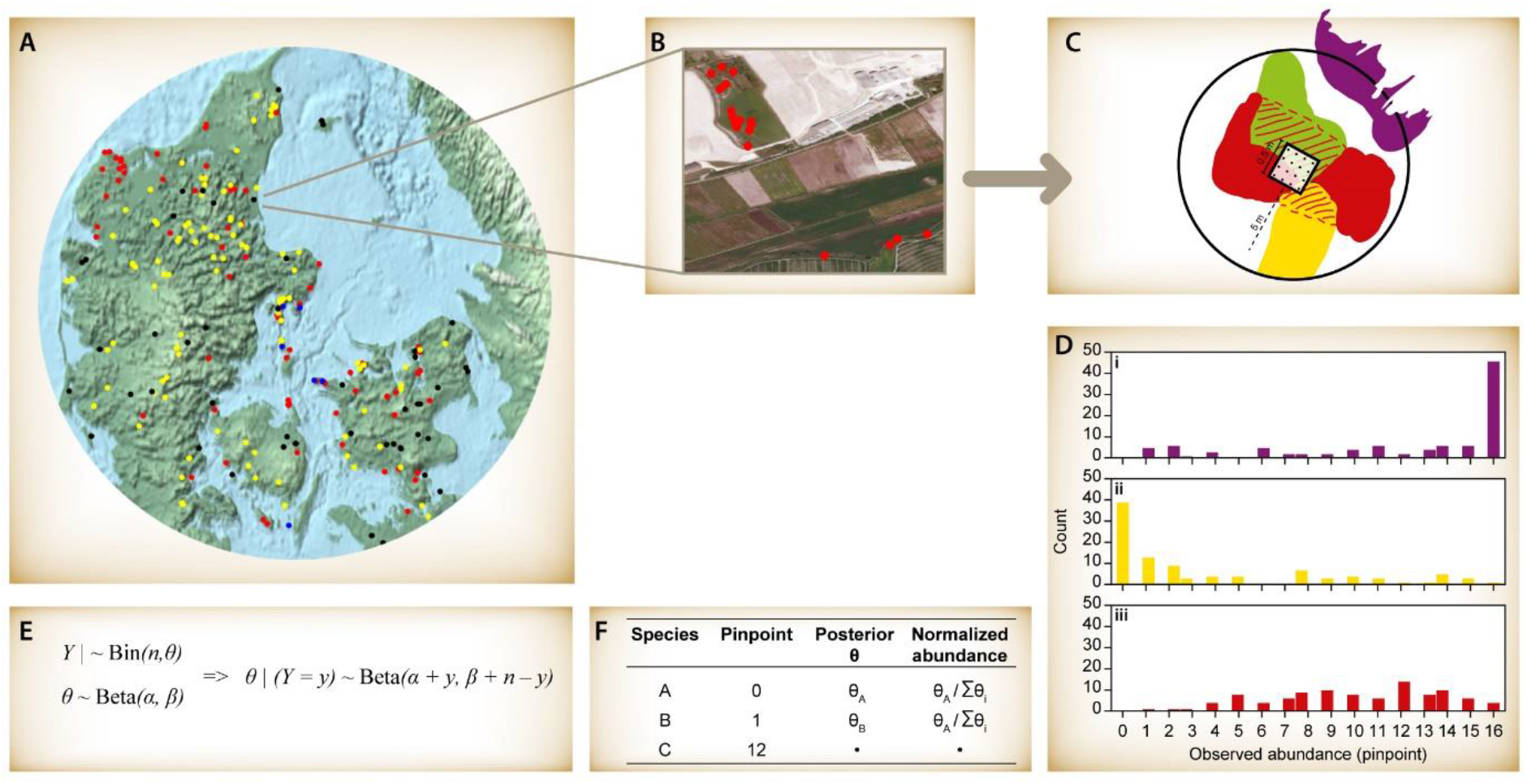
Conceptual figure. A: overview of NOVANA with sites distributed in different NATURA 2000 habitats. The different colors denote different habitat types. B: a local site consisting of several circles of 5m radius. C: within a circle, a pin-point frame with 16 grid points are used to measure cover. The three colors denote different plant species. D: empirical distribution of cover of three species across all sites. These empirical distributions are used as priors for each species local abundance. E: Bayesian framework to obtain updated (posterior) local abundance. F: example of observed cover and posterior (updated) relative abundance.

Generally, the two-stage sampling design allows for more than a single subsample in each large plot (e.g. Skinner 1986), but here we will only consider the case of a single subsample of plant abundance. While such a two-stage sampling design provides good local estimates of both the number of species and the relative abundances, it is still a problem to calculate Hill diversity indices at the local scale, because some of the species found in the large plot are not present in the small plot and such species are then incorrectly weighted with zero relative abundance.

The aim of this study is to suggest a method for calculating local Hill diversity indices from species richness and relative abundances data that is collected using the above-mentioned two-stage sampling design. We suggest to replace the locally estimated relatively abundances with Bayesian updated relative abundance estimates, where the prior probability distribution of the relative abundances are empirically estimated from all plots of the same habitat types. We assume that the relative abundance is measured by cover using the pin-point method, but the used Bayesian updating method may be applied on other plot-based measures of plant abundance.

## Methods

### Bayesian updated Hill diversity

Most plant species are spatially aggregated due to vegetative growth and limited seed dispersal (Watt, 1947, Pacala & Levin, 1997), and therefore plot-based estimates of plant abundance data typically displays L-shaped or U-shaped distributions, which may adequately be modelled using the beta distribution (Damgaard and Irvine 2019). Furthermore, when plant cover is sampled using the pin-point method it is recommended to model the distribution of pins that touch a specific species by the beta-binomial mixture distribution (Damgaard and Irvine 2019).

When working with Bayesian inference, the parameters in the prior probability distribution of all parameters underlying the sampling process has to be specified. This can be difficult without substantial prior empirical information and an uninformative prior is often chosen. Here, we propose a Bayesian framework that borrows strength from this design where variation in the abundance of a focal species can be learned from variation across samples. More precisely, we adopt an empirical Bayes approach (Carlin and Louis 1996) and rely on a prior probability distribution that reflects the relative species abundance in all other plots of the same habitat type. For each species, the prior probability distribution was assumed to be a beta distribution, which was fitted to the sampled cover from plots that was classified to belong to the same habitat type.

Following the logic of Bayesian inference, we use the likelihood function of the beta-binomial distributed locally observed pin-point cover data to update our prior beta distribution of the cover. We model the likelihood of the observed pinpoint data *P*(*Y* = *y_i_*) for species *i*, as a binomial distribution *Bin*(*n, q_i_*), where *n* is the number of grid points in the pin-point frame (the maximum number of possible hits) and *q_i_* is the local cover of species *i*. The empirical prior distribution of the local cover is assumed to be beta distributed as *q_i_*~*Beta*(*α_i_, β_i_*), where *α_i_* and *β_i_* are estimates obtained from the empirical distribution of the observed cover across all plots.

The resulting posterior probability distribution of the local cover is then the conjugate beta distribution,

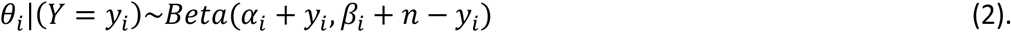

The mean of the posterior probability distribution, 0 < *E*(*θ_i_*) < 1, is then used as an estimator of the local cover of plant species *i*, and we may calculate a Bayesian updated estimate of the local relative abundance,

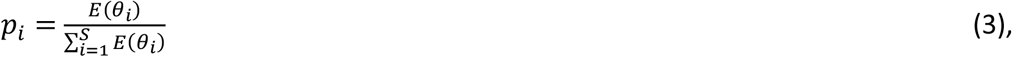

which are larger than zero for all *S* species, i.e. also for plant species that are observed in the large plot but not observed in the small plot. We then use these Bayesian updated estimates of the local relative abundance to calculate the local Hill diversity indices.

### Posterior predictive checks

The use of the Bayesian updated estimator was validated using posterior predictive checks (Gelman et al. 2003). That is, we drew a sample from the posterior probability distribution of species abundance for each species in a plot, and denote these the generated cover data. We repeat this process 1000 times, so that we get a distribution of generated cover data for each plot. If the model assumptions are appropriate the generated cover data will resemble the observed cover data when viewed through a relevant test statistic. The test statistic used here is the calculated Hill-Shannon diversities.

### Case study: Plant cover data

In the Danish monitoring program NOVANA, the absence-presence data of all higher plant species at a site is estimated in ten randomly positioned circles with 5-meter radius, and at the center of this circle plant cover data was estimated in 0.5m x 0.5m quadrates with the pin-point method using a horizontal frame with a 4×4 grid with the 16 intersections at a distance of 10 cm. At each intersection, a sharply pointed pin with a diameter of 0.5 mm was passed vertically through the vegetation and the cover of a species is measured by the proportion of the inserted pins that touches the species (Nielsen et al. 2012).

Only abundance data (sampled in the small plots) from plots where the species was observed in the large plot was used in the fitting of the prior probability distribution.

Using a subset of the collected monitoring data, the Bayesian updated Hill-Shannon diversity index was calculated for plots that were classified as Nardus grasslands (EU 2013) and sampled in 2014 (Nielsen et al. 2012). Subsequently, the calculated Hill-Shannon diversity indices were plotted against the soil pH measured at the sites (Nielsen et al. 2012).

### Software

Software for calculating the Bayesian updated Hill diversity indices, as well as workout examples may be found in the Electronic Supplement.

## Results

Both the observed species richness and the calculated Bayesian updated Hill-Shannon diversity indices in Nardus grasslands both increased with soil pH (Fig. 2). This positive effect of soil pH on species richness and diversity is expected, and similar results has been found in other studies and habitat types (Pärtel 2002).

**Fig. 2.**
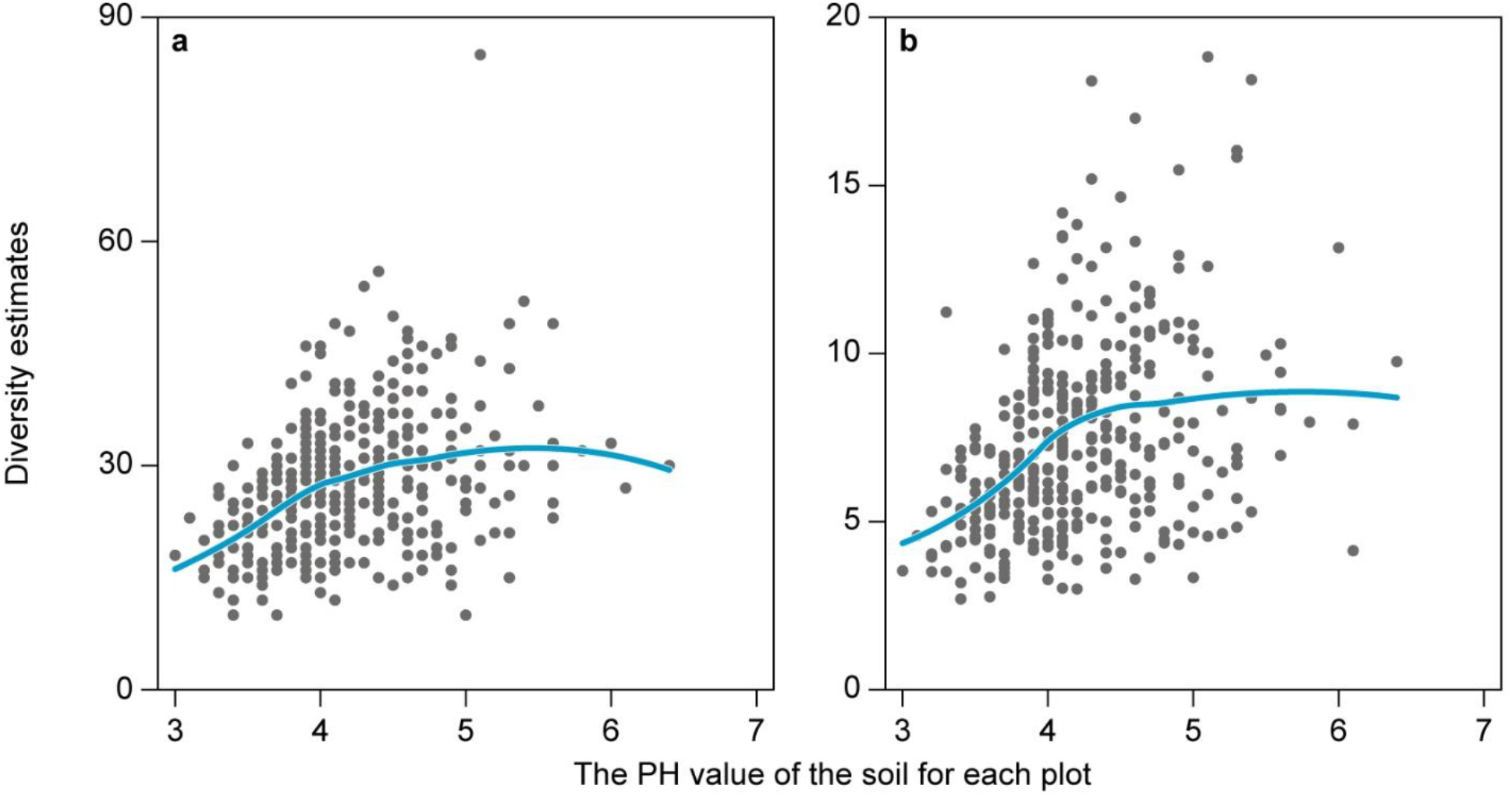
Species richness (a) and Hill-Shannon diversity (b) in Danish Nardus grasslands plotted against the measured soil pH at the site. The blue lines are the fitted smoothed conditional means using the loess method.

To validate the Bayesian updated estimator of relative abundance, the mean of the posterior generated Hill-Shannon diversities for each acid grass land was plotted against the calculated Hill-Shannon diversity for the same plot (Fig. 3). Eight percent of the plots fall outside the 95% credibility interval of the posterior predictive check, and these plots are mainly plots with relatively low diversity. This indicate that the Bayesian updated estimator of relative abundance may somewhat underestimate diversity at plots with relatively low diversity.

**Fig. 3.**
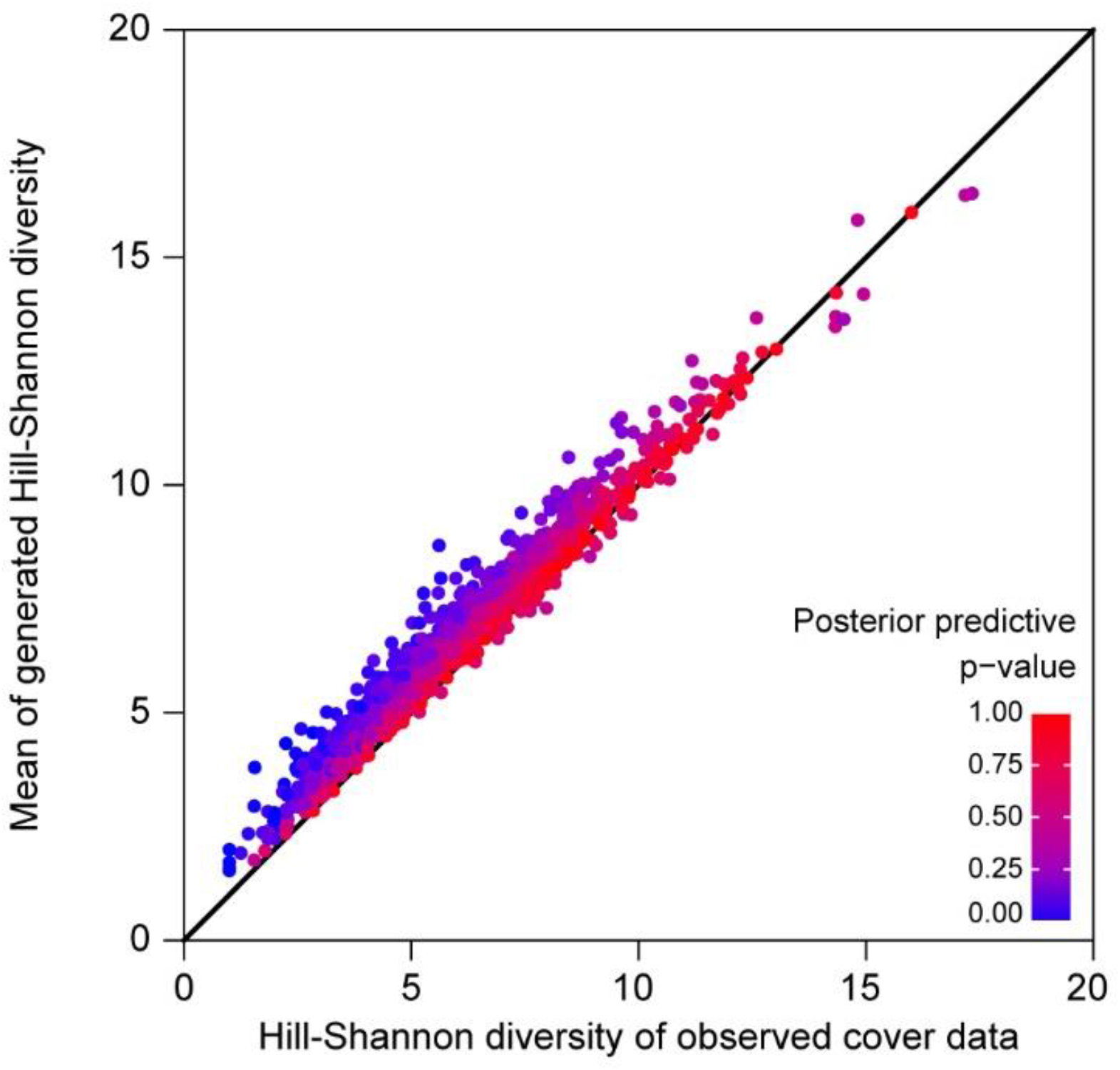
Mean of the posterior generated Hill Shannon diversities for each plot against the observed Hill Shannon diversity for the same plot. Each point in the scatter plot is colored according to the posterior predictive p-value of the plot.

## Discussion

Several authors have previously pointed out serious sampling issues when using species richness as a measure of diversity and have instead recommended to use Hill diversity indices where the number of species is weighted by their relative abundance (e.g. Haegeman et al. 2013; Roswell et al. 2021). Here, we complement this recommendation by the suggestion to use a two-stage sampling design for measuring plant species diversity. Such a sampling design will generally permit good estimates of both plant species richness and abundance. However, in order for this suggestion to be operationally for calculating Hill diversity indices, a positive estimator of the relative abundance is needed, and here we have suggested a Bayesian updated estimator of relative abundance, which is always larger than zero. In the performed case study on Nardus grasslands the Bayesian updated Hill-Shannon indices were easy to calculate and behaved as expected, although posterior predictive checks suggested that the index may be biased when diversity is low.

The use of the mean Bayesian posterior probability as a suitable and robust estimator has previously been advocated to treat zero-values at low sample sizes in ecological studies (Damgaard and Fayolle 2011).

The suggested two-stage sampling design is a special case of a more general class of two- or multi-stage sampling designs also known as cluster sampling designs, where the typical objective is to stratify the observed variation in abundance into primary units from which you may take a random sample.

Here we have only considered plant species diversity but the principle may easily be generalized to other domains where it is relevant to calculate diversity indices, e.g. molecular ecology methods relying for instance on 16S amplicon sequencing to probe the diversity and abundance of different taxa in a community.

## Supporting information

R function

## Acknowledgements

The main results of this study were obtained during a student project course organized by Asger Hobolth.

## Notes

### Competing Interest Statement

The authors have declared no competing interest.

